# Tissue signatures of human macrophages during homeostasis and activation

**DOI:** 10.1101/2025.05.23.655632

**Authors:** Daniel P. Caron, William L. Specht, David Chen, Steven B. Wells, Peter A. Szabo, Peter A. Sims, Donna L. Farber

## Abstract

Human macrophages (MΦs) reside in tissues and develop tissue-specific identities. While studies in mice have identified molecular signatures for site-specific MΦ differentiation, we know less about the transcriptional profiles of human MΦs in distinct sites, including mucosal tissues and lymphoid organs during homeostasis and activation. Here, we use multimodal single-cell sequencing and *ex vivo* stimulation assays to define tissue signatures for populations of human MΦs isolated from lungs, small intestine, spleen, bone marrow, and lymph nodes obtained from individual organ donors. Our results reveal distinct tissue-adapted gene and protein profiles of metabolic, adhesion, and immune interaction pathways, which are specific to MΦs and not monocytes isolated from the same sites. These signatures exhibit homology to murine MΦs from the same sites. Tissue-adapted MΦs remained responsive to polarizing cytokine stimuli *ex vivo*, with upregulation of expected transcripts and secreted proteins, while retaining tissue-specific profiles. Together, our findings show how human MΦ identity is coupled to their site of residence for mucosal and lymphoid organs and is intrinsically maintained during activation and polarization.

## INTRODUCTION

Macrophages (MΦs) are myeloid-lineage cells that populate diverse tissues and play critical roles coordinating tissue homeostasis, mediating innate responses to pathogens, promoting adaptive immunity, and facilitating resolution of inflammation^1^. MΦ populations initially develop from non-hematopoietic precursors during fetal development and are maintained throughout life by self-renewal or replenishment from monocytes^2, 3^. Importantly, MΦs adopt tissue-specific profiles; studies of murine MΦs across sites have uncovered site- and niche-specific gene expression patterns, comprising distinct regulatory mechanisms, and corresponding functional profiles^4–10^. These site-specific features are less clearly defined in humans, but MΦs are implicated in tissue-specific anti-pathogen responses, protective immunity, and disease pathologies such as non-alcoholic steatohepatitis (NASH) in liver, acute respiratory distress syndrome (ARDS) in the lungs, and atherosclerosis in blood vessels^11–13^. Defining their specific roles in both healthy and diseased contexts will require a greater understanding of how MΦs adapt to human tissues during homeostasis and activation.

As obligate tissue residents, MΦs are programmed by their local microenvironment and exhibit tissue-specific adaptations based on phenotype and function^4, 9, 14^. Lung alveolar MΦs respond to local signals by upregulating molecules necessary for uptake, transport, or catabolism of surfactant lipid components (e.g., phospholipids and cholesterol)^5, 15^, while splenic red pulp MΦs respond to local heme by upregulating hemolysis programs, including enzymes involved in releasing heme-associated iron and degrading byproducts^16^. In addition to modulating homeostatic and metabolic pathways, tissue-specific adaptations in MΦs can also directly regulate immune-related responses. For example, molecular programs in intestinal lamina propria MΦs induce an immunotolerant phenotype, suppressing inflammatory responses^17^ and encouraging expansion of tolerogenic regulatory T cell populations^7^. In tissues where MΦs are less abundant (such as lymph nodes or bone marrow), profiles of MΦ tissue-specificity and heterogeneity are limited. While tissue-specific programs are relatively well-characterized in murine models^4, 8, 10^ and MΦs profiled from human disease explants indicate core functions may be conserved across species^18–21^, evaluations of these tissue-resident cells’ diverse roles within a healthy human context has remained elusive.

As highly responsive immune sentinels, a key feature of MΦ function is their capacity to assume new phenotypic identities in response to environmental signals^22, 23^. Although MΦ plasticity exists on a complex spectrum^24^, *ex vivo* treatment paradigms have been established to evaluate MΦ polarization to various pro-inflammatory (M1-like) or anti-inflammatory (M2-like) environmental cues^22, 24^. In murine models, tissue resident MΦ populations can respond to stimuli with distinct kinetics, transcriptional regulation, and downstream functional responses depending on site of origin^25^. Unfortunately, our understanding of how human MΦs respond to these and other stimuli is largely limited to studies of MΦs derived *in vitro* from monocytes or stem cells, not tissue-resident MΦs^24, 26, 27^. Thus, it is unclear whether tissue-specific molecular programs in human MΦs affect their responsiveness to stimuli or vice versa. Using the paradigm of M1-like and M2-like cues, we evaluate the degree to which MΦ tissue-specification affects responsiveness to stimuli, and how this functional plasticity affects maintenance of tissue adaptations.

Recent efforts for transcriptomic profiling of human MΦs have relied on integration of scRNA-seq from diverse disease states, protocols, tissue-sources, or species^28, 29^. While these studies have provided important context for MΦ function in disease, effective investigation of tissue-specific profiles at homeostasis requires comparison between multiple sites within the same individual. Here, we use a well-validated organ donor resource^30^ to investigate MΦs and monocytes from six distinct sites (blood, BLD; lung, LNG; jejunum lamina propria, JLP; spleen, SPL; bone marrow, BMA; and lymph nodes, LN), where they play key roles in lymphoid and mucosal immunity and homeostasis^2, 31, 32^. First, we analyzed monocytes and MΦs from a previously generated human immune cell atlas^33^, identified canonical tissue MΦ subsets, and extracted tissue-specific gene and protein signatures. Then, we evaluated the stability of these tissue-signatures and functional plasticity of tissue-specified MΦs from new donors in *ex vivo* culture system—either at rest or under M1/M2 polarizing conditions with bacterial lipopolysaccharide (LPS) and interferon-γ (IFNγ) or interleukin 4 (IL-4). Together, these studies reveal how human tissue impacts these innate immune cells at homeostasis.

## MATERIALS AND METHODS

### Sample acquisition

Blood (BLD), lung (LNG), jejunum lamina propria (JLP), and spleen (SPL) samples for *ex vivo* experiments were obtained from deceased human organ donors^30, 34^. The use of organ donor tissues is not considered human subjects research as confirmed by the Columbia University IRB because the donors are deceased. Demographic information of donors used in this study is listed in Table S1.

### Ex vivo macrophage and monocyte cultures

Mononuclear cells from each tissue-site were isolated by mechanical and enzymatic digestion followed by density centrifugation, as previously described^35–39^. For BLD, LNG, and SPL samples, further erythrocyte depletion was accomplished by a brief (2min) incubation in ACK lysis buffer (Gibco) at 37°C. To deplete epithelial cells from the JLP, samples were incubated for 5 min in TruStain FcX blocking buffer (BioLegend), stained for 10 min with biotinylated α-CD326, and subjected to negative selection by streptavidin and annexin V microbubbles following manufacturer’s instructions (Akadeum). Samples were washed with buffer (DPBS, 5% FBS, 2mM EDTA) by centrifugation (400 x g, 5 min, 4°C), resuspended in Zombie NIR Viability dye (BioLegend) for 20 min, washed twice, incubated for 10 min in TruStain FcX and Monocyte Blocker (BioLegend) and stained with surface antibodies for 20 min (Table S3). Samples were sorted using a FACS Aria II (BD Biosciences) to isolate monocytes and MΦs (CD45^+^ CD33^+^ CD3^-^ CD19^-^ CD66b^-^ CD117^-^ CD326^-^ cells; Fig. S4A, Table S2).

Cells were seeded at 100,000 cells per well and cultured for 12 hours at 37°C in 0.5 mL media (RPMI, 10% FBS, 1X PSQ) on 24-well Nunc UpCell plates (ThermoFisher) either untreated, or treated with 0.02μg/mL LPS (Sigma) and 40ng/mL recombinant human IFNγ (R&D Systems), or 20ng/mL recombinant human IL-4 (R&D Systems). UpCell plates were incubated on ice for 20 min for non-enzymatic cell dissociation. Culture supernatants were cryopreserved, and samples were stained for 30 min with TotalSeq-C Hashtags following manufacturer’s directions (BioLegend). Samples were counted, pooled at equal proportions, treated with human TruStain FcX and stained for 30 min with TotalSeq-C Universal Cocktail following manufacturer’s directions (BioLegend). Samples for each donor were loaded across two lanes of a 10X Genomics Chromium instrument targeting between 8,000-10,000 cells each. cDNA synthesis, amplification, and sequencing libraries were prepared using Next GEM Single Cell 5’ Kit v2 (10X Genomics) and sequenced on a NovaSeq X (Illumina) with 28 cycles for read 1 and 90 cycles for read 2.

### Tissue-immune cell atlas pre-processing

To derive tissue signatures, we extracted MMoCHi-annotated^40^ monocyte and MΦ events from a CITE-seq atlas of immune cells isolated from human organ donor tissue^35^. Donors or libraries without antibody-derived tag (ADT) profiling (donors 582C and 583B; library 637C_6) were removed. For downstream analysis, gene expression (GEX) and antibody-derived tag (ADT) counts were library size and log-normalized to log(counts per 10k+1) and log(counts per 1k+1), respectively. Landmark-registration, as previously described^35, 40^, was used to integrate ADT expression across donors (*mmochi* v0.2.1).

Tissue-specific embeddings were computed on MΦs from LNG (bronchoalveolar lavage and parenchyma samples merged), JLP, SPL, bone marrow (BMA), and lymph nodes (LN; both lung-associated and mesenteric), and a joint-embedding was computed for monocytes from blood (BLO) and all tissue sites. Non-protein-coding, mitochondrial, ribosomal, or hemoglobin genes, and immune-receptor ADTs were removed prior to embedding. Principal components were computed on the top 5000 most highly variable genes in each donor (*scanpy* v1.10.4), or on all ADTs with between 20-80 percent positivity across the sample, integrated by donor using Harmony^41^ (*harmonypy*; v0.0.9), and used to compute neighbor graphs (50 nearest; *scanpy*). Merged, multimodal neighbor graphs were created using weighted-nearest neighbors^42^, embedded using Uniform Manifold Approximation and Projection (UMAP) with a minimum distance between 0.5 and 0.65, and clustered using the Leiden algorithm^43^ at a resolution between 1 and 1.4 (*muon*; v0.1.7). Clusters were manually annotated by marker expression. Similarity of MΦ subsets isolated from the two LN sites was determined using harmony-integrated principal component embeddings computed for the merged LN samples.

### Pre-processing of ex vivo culture sequencing

Alignment to the GRCh38 reference (Gencode v24 annotation) was performed using Cell Ranger (v7.1.0; 10X Genomics). Events with under 250 unique genes profiled, 1,000 GEX counts or 50 ADT counts, and events with over 50,000 GEX counts, 10,000 ADT counts, or 20% mitochondrial counts were removed. Hashtags were demultiplexed using HashSolo^44^ with priors determined by commercially provided multiplet rates (*scanpy*). HashSolo-inferred multiplets and hashtag-negative events were removed. An initial clustering on highly variable genes (as above) for each donor was used to identify and remove clusters enriched for markers of contaminant lineages (e.g., *G0S2*, *CD3E*, *CD19*, *JCHAIN*, *KRT7*, *DCN*; <2% of total events) or low-quality events (abnormally high mitochondrial content or low GEX counts). Next, clustering and marker-based manual annotation was performed for each tissue/treatment combination on the library-integrated (*harmonypy*) principal components of the GEX, using a Leiden resolution of 2.5 (*scanpy*). A global, library-integrated UMAP embedding was computed for visualization.

### Pseudobulk differential gene and protein expression analysis

GEX counts were aggregated across each sample or each sample/subset combination. Samples with under 50 (for the immune atlas) or 20 (for the *ex vivo* stimulation) cells, or genes not represented in 15% of samples with at least 20 counts were excluded (*dreamlet* v1.4.1). Differential expression (DE) was estimated using linear mixed models^45^ with either a random intercept of donor or—to derive universal stimulation signatures—random intercepts of donor and tissue. Differential percent-positivity analysis for ADT expression was performed with analogous mixed models (*statsmodels*; v0.14.0). One-v-one and one-v-all effects were computed using contrasts. P-values were adjusted for false discovery rate (Benjamini-Hochberg method). Full differential expression tables are included in Table S3.

Gene signatures associated with specific tissues or stimuli were extracted as the top 100 significantly upregulated (p-adj<0.05, log_2_(fold-change)>0) protein-coding, non-ribosomal genes by t-statistic. Signature scores for each cell were calculated using *scanpy*. MΦs in cell cycle were quantified using *scanpy* and gene lists associated with S or G2/M phase^46^. Gene Set Enrichment Analysis (GSEA) and gene ontology (GO) were performed by *gseapy* (v1.1.3). Enrichment of tissue-signatures in the estimated log_2_(fold-change) between pairs of tissues or subsets was evaluated by pre-ranked GSEA. Enrichment of these signatures in the Tabula Sapiens^47^ was evaluated by pseudobulk DE across donor/tissue combinations with at least 10 MΦs (*scanpy*). Enrichment of these signature in murine tissues was evaluated using the fold change in expression in bulk RNA-seq data generated previously on sorted MΦs^4^.

### Cytokine analysis

Supernatants were shipped to Eve Technologies Corp. (Calgary, Alberta) to apply a bead-based multiplexed quantification of 71 human cytokines and chemokines. For each analyte, concentrations were derived by aligning beads’ fluorescence intensity to standard curves and multiplying by the dilution factor (1.8x). For statistical analysis and visualization, values were clipped to the provided limit of detection. Enrichment or depletion in a particular tissue-site upon stimulation was determined by a paired t-test on log_10_-transformed concentration.

## RESULTS

### Heterogeneous macrophage populations within tissues

To identify underlying heterogeneity within monocyte and MΦ populations from blood (BLD) and five tissue-sites including lungs (LNG), jejunum lamina propria (JLP), spleen (SPL), bone marrow (BMA) and lymph nodes (LN—both lung-associated [LLN] and mesenteric [MLN]), we extracted over 200k MΦ and monocyte events from a recent CITE-seq atlas of immune cells from 24 human organ donors’ immune cells^35^ and performed sub-clustering (Figs. 1A, S1, and S2, Table S1). We identified major subsets of monocytes, including classical (*CD14*^hi^ *S100A8*^hi^ *FCN1*^+^ *VCAN*^+^) and non-classical (*FCN1*^+^ *VCAN*^mid^ *FCGR3A*^+^ *C1QA*^mid^) populations^28^, which were distributed across tissue-sites (Figs. 1A-B, S1B-D). In contrast, we uncovered heterogeneity within each tissue’s MΦ compartment, consistent with previously described site- and/or niche-specific populations. The lungs contained interstitial (*LGMN*^+^ *MARCKS*^+^)^48, 49^ and alveolar (*PPARG*^+^ *FABP4*^+^ CD11b^-^ CD11c^+^)^49, 50^ MΦs, while the intestines were predominantly composed of mucosal MΦs (*RUNX3*^+^ HLA-DR^hi^ *CD209*^+^)^31, 51^, and the spleen contained red pulp MΦs (*CD14*^-^ *CD16*^+^ *SPIC*^+^ *VCAM1*^+^)^6, 32^ (Figs. 1B, S1E-G, S2A-C). In the BMA, we identified vascular-associated MΦs consistent with previously described osteal MΦ populations (*LYVE1*^+^ HLA-DR^lo^ CD11b^+^ CD163^+^)^52–54^ and a small population of progenitor niche MΦs (*SPIC*^+/-^ *VCAM1*^+/-^ CXCR3^+^)^55, 56^ (Figs. 1B, S1H, S2D). Across both LN sites, we identified T cell zone (*CD68*^+^ CD169^-^)^57, 58^ and subcapsular/medullary sinus (*CD68*^+/-^ CD169^+^)^57–59^ MΦs (Figs. 1B, S1I-J, S2E-F). Due to the high similarity in transcriptomes and surface proteomes between both sites (Fig. S2G-H), LN samples were merged for further analysis.

**Figure 1:**
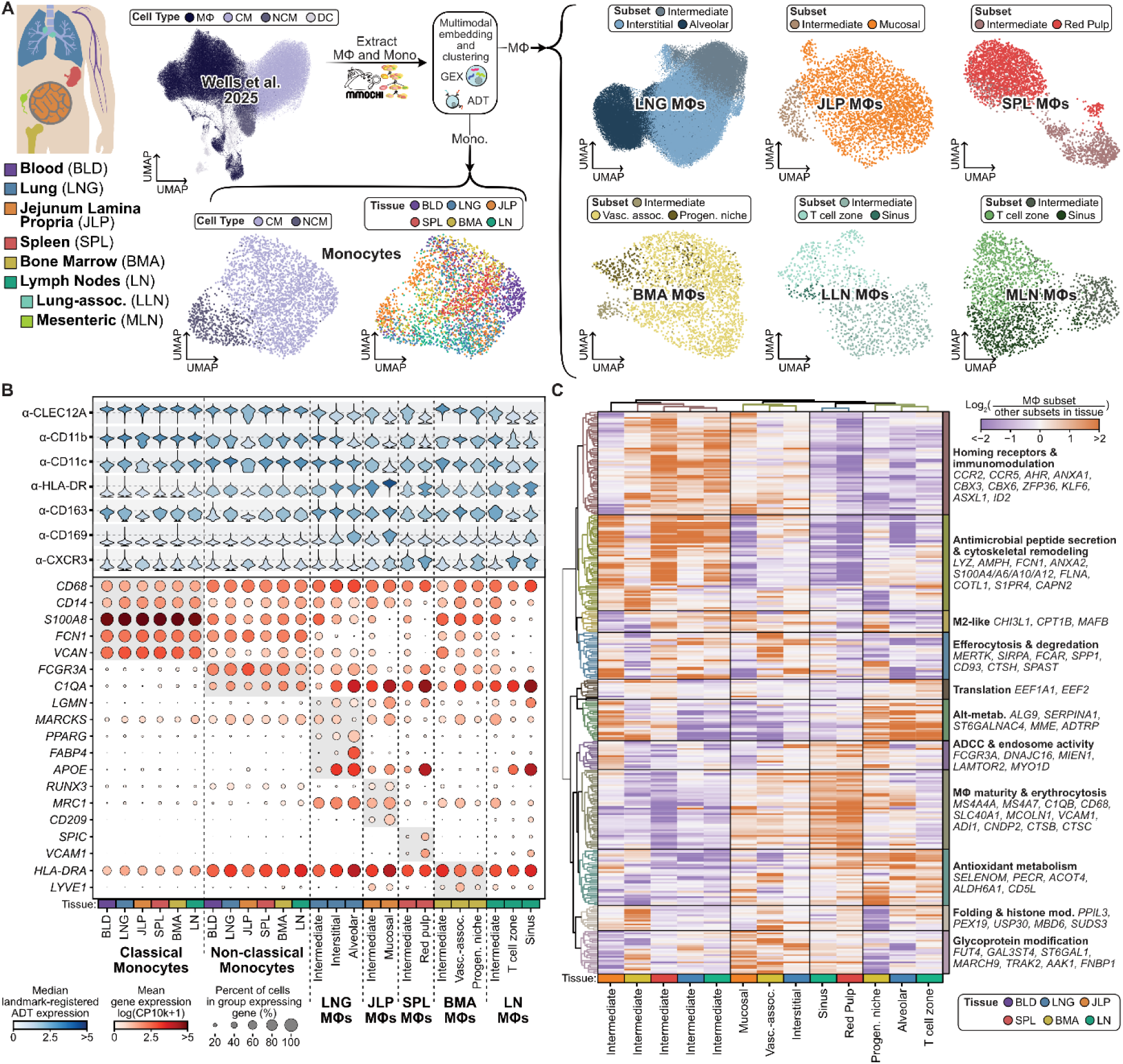
Diverse populations of macrophages exist within human tissues. **(A)** Schematic of analysis strategy. Macrophages (MΦs) and monocytes were extracted from an organ donor atlas of tissue immune cells. MΦs from each tissue and monocytes were subclustered and manually annotated using antibody-derived tag (ADT) and gene expression (GEX). Uniform manifold approximation & projection (UMAP) plots are colored by classified cell type, tissue-of-origin, or manually annotated MΦ subset. **(B)** Dot and violin plots display GEX and ADT expression of selected subset markers. Dots are colored by mean GEX and size reflects the percent of cells expressing a gene. Violins display the distribution of landmark-registered ADT expression and are colored by the median. **(C)** Heatmap of log_2_(fold-change) of top 25 differentially expressed genes for each subset when compared to other MΦ subsets from the same tissue. Rows and columns are clustered by Spearman correlation.

In all tissues examined, we also detected a population that exhibited features of an intermediate MΦ subset, marked by shared expression of monocytic and MΦ markers (*S100A8*^mid^ *FCN1*^mid^ *VCAN*^mid^ *C1QA*^mid^) (Figs. 1B, S1K), suggestive of early stages of MΦ differentiation^48, 51, 60^. Intermediate MΦs were relatively frequent, representing between 10-50% of MΦs in each tissue (Fig. S1K). Analysis of differentially expressed genes between MΦ subsets within each tissue revealed similar profiles of intermediate MΦs across tissues, with increased expression of homing receptors (*CCR2*, *CCR5*) and cytoskeletal remodeling (*FLNA*, *COTL1*, etc.) in intermediate MΦs compared to their more tissue-specified counterparts (Fig. 1C, Table S3). Differential expression also revealed other MΦ subsets with functional overlap, including LN sinus MΦs and splenic red pulp MΦs, which shared increased expression of markers of MΦ maturity (*MS4A4A*, *MS4A7*)^61^ and erythro-efferocytosis (*VCAM1*, *SLC40A1*, *ADI1*) (Fig. 1C). In contrast, LNG interstitial MΦs, JLP mucosal MΦs, and BMA vasculature-associated MΦs shared genes associated with M2-differentiation (*CHI3L1*, *MAFB*)^62^ and glycoprotein modification (*FUT4*, *GAL3ST4*, etc.), while LNG alveolar MΦs, LN T cell zone MΦs, and BMA progenitor-niche MΦs shared upregulation of lipid metabolism pathways (*CD5L*, *ACOT4*, *PECR*) (Fig 1C, Table S3). Additionally, across tissues and subsets, a fraction of MΦs were captured in various stages of cell cycle (see Methods; Fig. S1L-N). Together, these results support recent findings that despite niche-specific identification, common functional requirements may drive conserved molecular programs in MΦ subsets across tissues^28, 63, 64^.

### Tissue-adaptations in macrophages, but not monocytes, drive unique molecular profiles

Having characterized diverse MΦ subsets within each tissue site, we next applied differential expression to identify molecular profiles associated with each site and investigate the extent to which different MΦ and monocyte subsets are tissue-adapted. We found unique transcriptional signatures associated with MΦs from each tissue site (Figs. 2A-B, S3A, Table S3). LNG and JLP MΦs had the highest number of differentially expressed genes (DEGs) across sites, while DEGs associated with LN MΦs were largely shared with SPL MΦs (Fig. S3B, Tables S3– 4).

**Figure 2:**
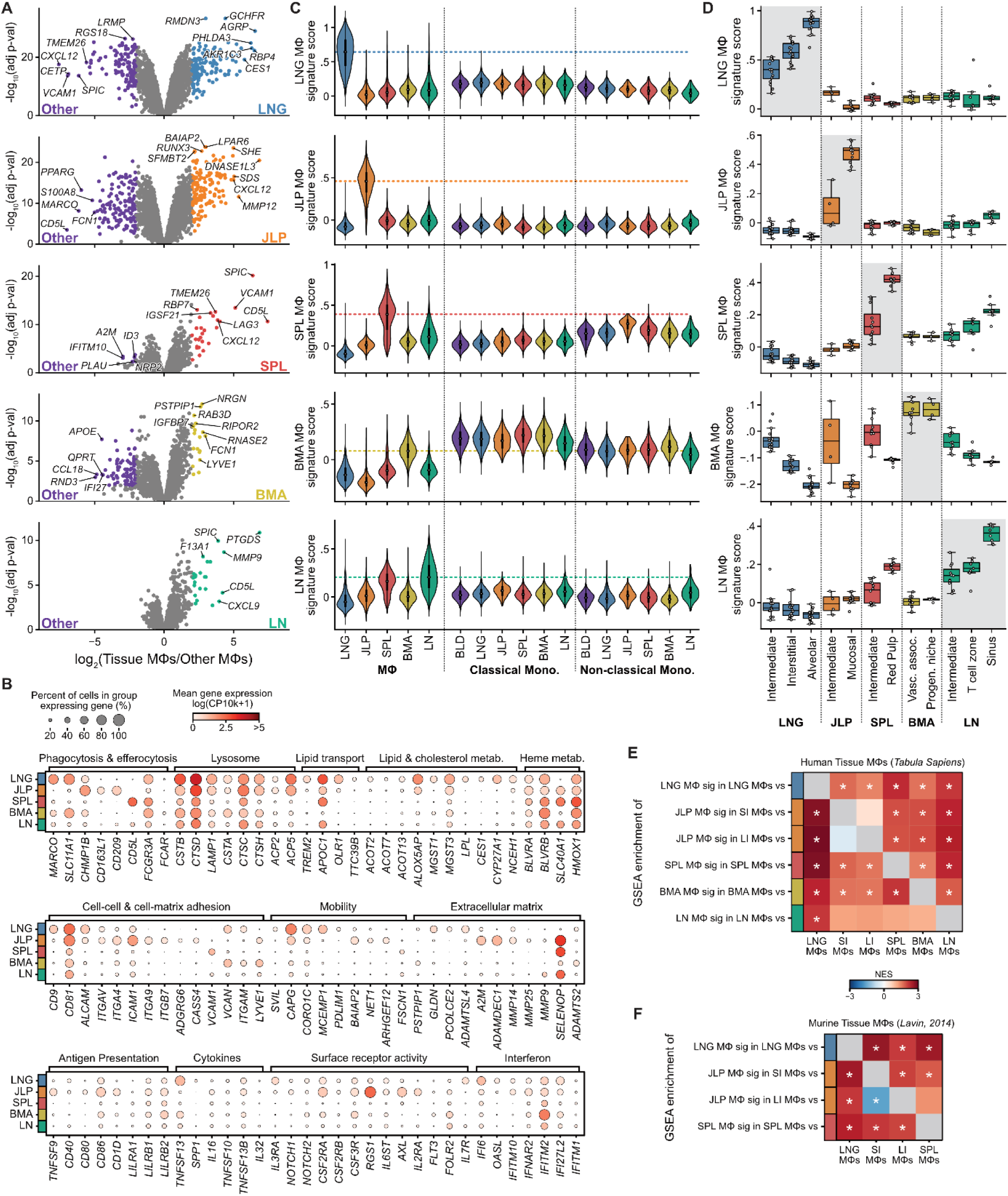
Tissue-specific adaptations in human macrophages drive distinct profiles. **(A)** Volcano plots highlight the top significantly differentially expressed genes (DEGs; adjusted p-value < 0.01 and |log_2_(fold-change)| > 2) between macrophages (MΦs) in each tissue site and all other tissue sites. DEGs are highlighted in purple (downregulated) or a tissue-specific color (upregulated). **(B)** Dot plots of selected DEGs, where dots are colored by mean GEX and size reflects the percent of cells expressing a gene. **(C)** Violins display expression scores of tissue-specific gene signatures (defined as the 100 most significant DEGs with log_2_(fold-change) > 1) for MΦs and monocytes in each site. **(D)** Boxplot displaying the mean tissue-signature score in each donor for each MΦ subset. Donors with fewer than 25 events in a tissue/subset combination were excluded. **(E-F)** Heatmaps displaying normalized enrichment score (NES) for tissue gene signatures (“Gene set”, denoted by color) in pairwise differential expression between tissue MΦs from the *Tabula Sapiens* **(E)** Lavin et al **(F)** dataset when MΦs from the same or similar tissue-sites (rows) are compared to other MΦ populations (columns). Asterisks denote significance (FDR adjusted p-value < 0.05) as assessed by pre-ranked gene set enrichment analysis (GSEA). LNG, lung; JLP, jejunum lamina propria; SPL, spleen; BMA, bone marrow; LN, lymph node; SI, Small Intestine; LI, Large Intestine

Across tissues, there were unique profiles of phagocytosis and related metabolism pathways, with specific phagocytosis receptors in the LNG (*MARCO*) and JLP (*CD163L1* and *CD209*, associated with anti-inflammatory resolution^65, 66^), as well as expression of the low affinity IgG receptor (*FCGR3A*) in MΦs from all sites except JLP (Fig. 2B). Concurrently, we observed tissue-specific expression of lysosomal and metabolic pathways, including high lysosomal protease (*CTSD*, *CTSH*) and inhibitor (*CSTB*, *CSTA*) expression in all sites except SPL, and enrichment of phosphatases (*ACP2* and *ACP5*) in LNG and JLP (Fig. 2B). Enrichment in LNG MΦs for lipid and cholesterol metabolism genes (*TREM2*, *APOC1*, *OLR1*, *ACOT2*, *LPL*) and in SPL MΦs for heme metabolism (*BLVRB* and *SLC40A1*) support known roles for cycling of surfactant^5^ and erythrocytes^16^, respectively (Fig. 2B).

MΦs across tissues also expressed unique markers of adhesion and mobility, with LNG-specific expression of tetraspanins (*CD9*, *CD81*)^67^, JLP-specific integrins (*ITGAV*, *ITGA4*, *ITGA7* and *ITGB7*), SPL-specific vascular adhesion marker expression (*VCAM1*) and BMA-specific enrichment for hyaluronic acid-binding molecules (*VCAN*, *LYVE1*), essential for lymphatic trafficking^68^ (Fig. 2B). LNG and JLP MΦs additionally expressed markers associated with increased mobility (e.g., *SVIL* and *CAPG* in the LNG; *NET1* and *ARHGEF12* in the JLP)^69–72^, and markers of extracellular matrix reorganization (*GLDN* and *PCOLCE2* in the LNG; *ADAMDEC1*, *MMP14*, and *MMP25* in the JLP) (Fig. 2B). Interestingly, *SELENOP*, a selenium transporter implicated in extracellular matrix regulation and migratory potential^73, 74^, was highly expressed in MΦs of the JLP, SPL, and LN (Fig. 2B).

Finally, we identified tissue-specific expression of key markers associated with immune interactions, including a mix of pro- and anti-inflammatory markers, primarily in JLP and BMA MΦs (Fig. S3A, Table S3). JLP MΦs additionally expressed high levels of antigen presentation and co-stimulation markers (*TNFSF9*, *CD80*, *CD86*, and *CD1D*) (Fig. 2B). LNG, BMA, and LN MΦ produced high levels of *TNFSF13* (APRIL) and *CCL18*, both associated with lymphocyte interactions, while JLP MΦs had high expression of the proinflammatory cytokine *IL16* and chemokines *CXCL12* and *CCL24* (Figs. 2B, S3A, Table S3).

Tissue-specific gene expression for surface receptors included high IL-3 receptor (*IL3RA*) in LNG MΦs, high Notch, GM-CSF, and G-CSF receptors (*NOTCH1*, *NOTCH2*, *CSF2RA, CSF2RB*, and *CSF3R*) in LNG, JLP, and BMA MΦs, and high folate receptor (*FOLR2*) in BMA and LN MΦs (Fig. 2B). Interferon response genes were also upregulated in various sites, including *IFI6* and *OASL* in LNG MΦs, and *IFITM2* and *IFI27L2* in BMA MΦs (Fig. 2B). Differential gene expression also uncovered tissue-specific expression of >20 transcriptional regulators, including *PPARG* in LNG MΦs, *RUNX3* in JLP MΦs, and *SPIC* in SPL MΦs, which align with transcriptional regulators identified for analogous subsets in mice^4, 15, 75, 76^ (Fig. S3A, Table S3). By profiling these cells using CITE-seq, we could additionally assess how surface protein expression changed across tissues (Fig. S3C). Protein expression reflected differentially expressed genes, including expression of CD123 (*IL3RA*) and LOX1 *(OLR1*) on LNG MΦs and expression of CD39 (*ENTPD1*), CD1d, and CD86 in JLP MΦs (Figs. S3C).

Most of these tissue signatures were uniquely expressed by the corresponding tissue’s MΦs compared to other MΦs, diminished in the corresponding populations of intermediate MΦs, and lowly expressed by tissue monocytes (Fig. 2C-D). The BMA signature however, composed of monocytic and vascular adhesion markers (e.g., *FCN1* and *LYVE1*), was highly expressed in BMA MΦs and across all monocyte subsets, but was not particularly enriched in BMA monocytes over other tissue sites (Fig. 2A-C). For the remaining tissues, these signatures were highly enriched in all MΦ subsets within a site when compared to MΦs derived from other tissue sites (Fig. S3D), albeit with variable expression between subsets (Fig. 2D). These results demonstrate our MΦ signatures derive from tissue-specific adaptations within cells, rather than from compositional differences across sites.

Finally, we assessed expression of these signatures in two external datasets: MΦs profiled as part of the *Tabula Sapiens*^47^ human tissue-atlas and MΦs sorted from murine tissues by Lavin and colleagues^4^, and found a high degree of concordance (Fig. 2E-F). Together, these results demonstrate that our MΦ signatures derive from tissue-specific adaptations within cells, rather than from compositional differences across sites, and uncovers many molecular markers that modulate tissue-specific functions in both immune regulation and homeostatic tissue maintenance.

### Macrophages respond to polarization stimuli while maintaining tissue-adaptation

Tissue-specific adaptations in MΦs could be maintained by cell-extrinsic (e.g., continuous microenvironmental signaling) or cell-intrinsic (e.g., epigenetic reprogramming) mechanisms^4, 77^. To dissect these possibilities, we examined the stability of tissue-specific adaptations *ex vivo* and evaluated whether MΦs in different sites exhibited tissue-specific responses to acute stimulation. MΦs and monocytes (CD45^+^ CD33^+^ CD66b^-^ CD117^-^ cells) were sorted from the BLD, LNG, SPL, and JLP, cultured for twelve hours with or without polarizing stimuli, including LPS+IFNγ for a pro-inflammatory M1-like stimulus and IL-4 representing an M2-like stimulus for tissue repair, and profiled by CITE-seq or cytokine array (Figs. 3A, S4A). Analysis of CITE-seq data revealed that cells segregated by stimulus condition (unstimulated, LPS+IFNγ, or IL-4) or cell type (monocyte or MΦ) (Figs. 3B, S4B-D). Differential expression identified global signatures of activation, in line with expected responses to LPS+IFNγ (e.g., *CIITA*, *IL27*, *GBP1*, *CXCL9*, *CXCL10*)^78^ or IL-4 (e.g., *CISH* and *MAOA*)^78, 79^, and activation of the downstream transcription factors (Fig. 3C-D, Table S3). These signatures were highly upregulated in both MΦs and monocytes across all tissues, suggesting conserved functional responses across sites (Fig. 3E,F).

**Figure 3:**
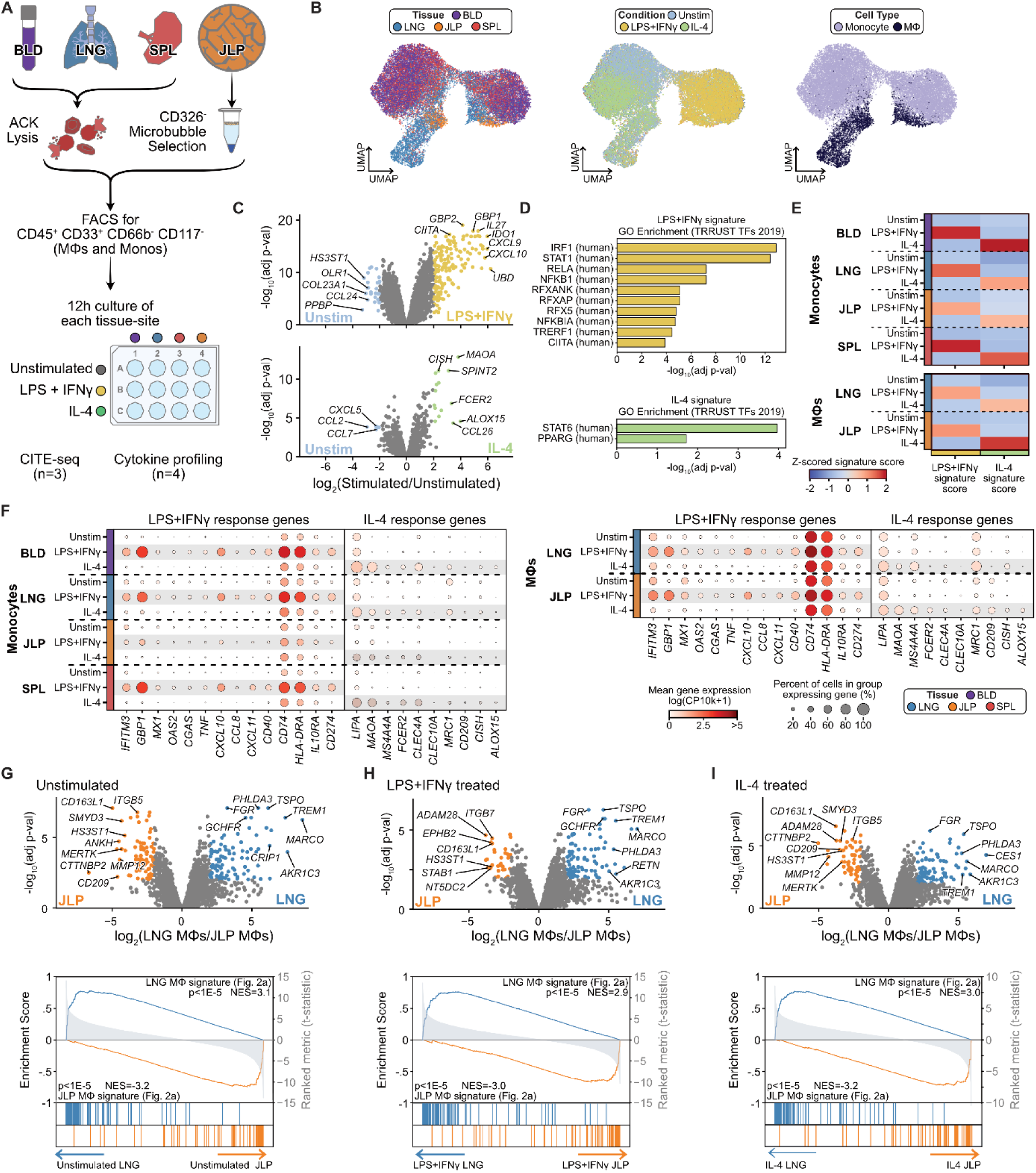
Macrophages maintain tissue identity during acute polarization. **(A)** Schematic of macrophage (MΦ) and monocyte enrichment and *ex vivo* culture. Blood (BLD), lung (LNG), and spleen (SPL) samples underwent ammonium-chloride-potassium (ACK) lysis, and jejunum lamina propria (JLP) samples underwent negative epithelial cell selection. Samples were sorted for CD45^+^ CD33^+^ CD66b^-^ CD117^-^ MΦs and monocytes, cultured for 12 hours either in untreated media, with LPS+IFNγ, or with IL-4 and subsequently harvested for CITE-seq and cytokine profiling. **(B)** Uniform manifold approximation & projection (UMAP) plots are colored by classified tissue-of-origin (“Tissue”), treatment condition (“Condition”), or annotated cell type (“Cell Type”). **(C)** Volcano plots display top significantly differentially expressed genes (DEGs; adjusted p-value < 0.01 and |log2(fold-change)| > 2) between treated and unstimulated cells. DEGs are highlighted in blue (downregulated upon stimulation), yellow (upregulated with LPS+IFNγ) or green (upregulated with IL-4). **(D)** Bar plot displaying significant overrepresentation of human transcription factor associated genes within condition-specific gene signatures (defined as the 100 most significant DEGs with log_2_(fold-change) > 1). **(E)** Heatmap of z-scored average stimulation signature scores for MΦs and monocytes in each site and condition. **(F)** Dot plots displaying expression of selected transcript markers of activation on monocytes (left) and MΦs (right) across tissues. Dots are colored by mean GEX and size reflects the percent of cells expressing a gene. **(G-I)** Volcano plots as in (C) displaying top DEGs between LNG and JLP MΦs in each condition. Below, pre-ranked gene set enrichment analysis (GSEA) plots display enrichment for LNG and JLP tissue-signatures (defined in Fig. 2A) within these differential gene expression comparisons. Genes are ordered left-to-right by their significance, raster-plots highlight genes present in the signature, and running average enrichment scores are shown by colored lines.

To evaluate whether the molecular tissue-signatures were conserved during *ex vivo* culture, we analyzed DEGs between MΦs from LNG and JLP, the sites with the highest numbers of MΦs profiled. Unstimulated MΦs maintained highly significant differential gene expression between the two sites, and these DEGs aligned with the signatures we had derived from MΦs isolated directly from tissue (Figs. 2A, 3G). These DEGs were maintained between LNG and JLP MΦs even upon activation with acute M1 or M2 polarization stimuli and were highly concordant with the previously derived tissue signatures (Fig. 3H-I), together suggesting that our tissue-specific profiles represent a separate axis of MΦ identity than acute polarization.

For a higher resolution assessment of the functional profile of *ex vivo* stimulated MΦs and monocytes, we performed multiplex cytokine profiling of culture supernatants. Among unstimulated samples, we detected high production of CCL2 and CXCL5 from the LNG and IL-16 from the JLP (Fig. S4F), confirming the tissue-specific patterns we previously detected at the transcript level (Figs. 2F, S2C). After stimulation with LPS+IFNγ, MΦs and monocytes across sites produced high levels of CXCL9 and CXCL10, consistent with upregulated expression at the transcript level (Figs. 4A-B, S4F). Additionally, we detected high induction of TNFα, CCL8, IL-1β, and IFNα2 in response to LPS+IFNγ in all cultures except JLP (Fig. 4A-B). After stimulation with IL-4, MΦs and monocytes across all samples produced increased CCL17 and VEGFA (Fig. 4C-D). IL-1RA was similarly induced in most cultures, but less prevalent in JLP (Fig. 4C-D). Secretion of inflammatory chemokines CCL2, CXCL1, and CXCL5 was reduced in IL-4-stimulated cultures (Fig. 4C-D). Together, these results indicate that despite diverse tissue-specific adaptations, MΦs maintain responsiveness to acute stimuli both in their transcriptome and, to a lesser extent, in their secretome.

**Figure 4:**
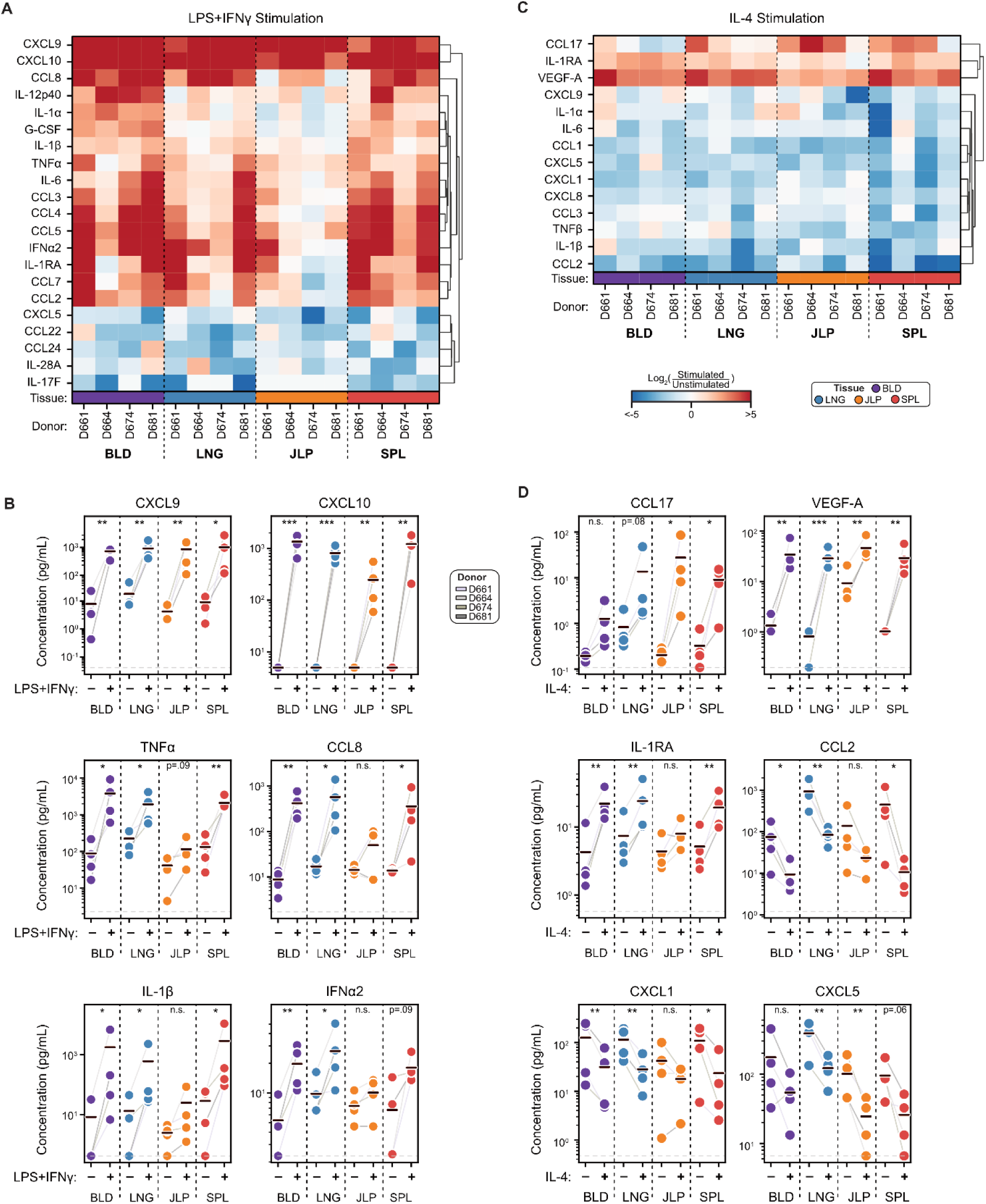
Cytokine production by *ex vivo* polarized monocytes and macrophages reveals reduced proinflammatory cytokines in intestine. Supernatants of unstimulated and stimulated samples were assessed by cytokine array. **(A)** Heatmap displaying the log_2_(fold-change) between LPS+IFNγ and unstimulated samples. Selected differentially produced cytokines and chemokines (|mean log_2_(fold-change)| >= 1) are hierarchically clustered. **(B)** Paired strip plots display concentration of selected cytokines in unstimulated (“-”) or LPS+IFNγ treated (“+”) samples. Means are indicated by black bars. Measurements from the same donor are connected by light gray lines. A dashed line displays the assay’s lower limit of detection. **(C)** Heatmap as in (A) displaying the log_2_(fold-change) between IL-4 and unstimulated samples. **(D)** Paired strip plots as in (C) for IL-4 treated samples. Significance is determined by pairwise t-test and indicated by ***, p < 0.001; **, p < 0.01; *, p < 0.05; or n.s., p > 0.1. For visualization, log_2_(fold-change) calculation, and significance testing, samples below the assay’s limit of detection were set to that value. LNG, lung; JLP, jejunum lamina propria; SPL, spleen; BMA, bone marrow; LN, lymph node

## DISCUSSION

In this study, we leveraged single-cell profiling to identify core tissue-adaptation signatures and niche-specific heterogeneity within human MΦs. These tissue-signatures are specific to MΦs, minimally expressed in their monocytic counterparts, and shared across intermediate MΦs and previously identified tissue-specific subsets. Tissue differences were maintained in the absence of tissue-specific cues (i.e. in *ex vivo* culture) and even during acute stimulation, suggesting that cellular tissue adaptation is controlled by cell-intrinsic transcriptional mechanisms. Together, our findings uncover human MΦ heterogeneity across diverse sites and provide reference signatures for their cellular states during homeostasis and following activation, important for designing therapies targeted to inflammatory responses in specific sites.

Because of the difficulty in obtaining viable MΦs from human tissues, studies evaluating tissue-specific profiles for human MΦs have required extensive integration or inclusion of diverse disease samples in order to obtain sufficient sampling for analysis^28, 47^. Here, we obtained tissue from organ donors, allowing for several improvements over previous work: first, our protocol enabled rapid isolation of live MΦs from freshly isolated samples, we obtained multiple sites from each individual, and applied both protein and transcriptome profiling along with a multimodal classifier for unambiguous annotation of MΦ populations distinct from monocytes, dendritic cells, and intermediate subsets. Using this approach, we significantly expanded upon tissue signatures previously identified in compiled human studies, and revealed additional tissue-specific profiles involving metabolic, activation, adhesion, and cell mobility pathways. Moreover, we identified high concordance between the transcriptional signatures we derived from humans with those established in murine MΦs^4^, despite vast differences in microanatomical structures^80, 81^, microbial burden^82^, and lifespan between species. Additionally, we detected tissue-specific enrichment of human orthologs to the TFs underlying murine tissue-signatures, such as *PPARG* in lung MΦs, *SPIC* in splenic MΦs, and *RUNX3* in intestinal MΦs^4, 15, 75, 76^, suggesting that murine models may provide insight into regulatory networks driving these molecular profiles in humans.

By profiling subset heterogeneity within the MΦ compartment of each tissue, we identified subsets that align with those described in previous human studies. Many of these subsets (e.g., alveolar and interstitial MΦs in the LNG^49^) were defined originally by niche-specific localization and later by functional markers. The multi-omic profiling performed here (with both surface proteome and transcriptome) may uncover new or improved markers for individual subsets. Additionally, we consistently identified populations of intermediate MΦs across tissues and captured cycling MΦs, suggestive of monocyte replenishment and self-renewal, respectively. The simultaneous occurrence of both of these processes in healthy tissue suggests that human tissue MΦ populations, in contrast to those in murine fate-mapping models^83, 84^, are not exclusively maintained by one process or the other. Instead, it supports a dynamic interplay between self-renewal and monocyte replenishment in population maintenance of tissue MΦs, as has been observed in human transplantation studies^51, 85, 86^.

The MΦ tissue signatures we describe were expressed in all MΦ subsets across niches within a tissue site, implying a tissue-wide mechanism of regulation. For example, we identified specialized phenotypes of MΦs within defined tissue niches, such as alveolar and interstitial lung MΦs, but also identified shared expression of tissue-wide transcriptional programs, suggesting that these subsets in humans either derive from common lung MΦ precursors and/or that their identifies are maintained by tissue-wide signals such as cytokines or interactions with structural cells. Notably, both intermediate and fully differentiated MΦ subsets exhibited tissue-specific signatures, while monocytes isolated from the same tissue did not. These results are analogous to our recent findings that tissue specific signatures were shared across multiple lymphocyte subsets (CD4^+^, CD8^+^, and γδ T cell subsets, natural killer cells, and innate lymphoid cells) but less pronounced in naive cells^35^, suggesting that differentiation may be required for immune cells to respond to tissue-specific cues.

When cultured *ex vivo*, devoid of tissue signals, lung- and intestine-derived MΦs preserved their tissue-specific molecular programs. These tissue differences persisted during acute (12h) polarizing signals comprising inflammatory (LPS+IFNγ) or repair-type (IL-4) stimulation. This maintenance of tissue-adaptations suggests that MΦs retain a robust memory of their tissue programming, which is not overridden by polarizing stimulation. One potential mechanism for this memory is epigenetic regulation of tissue-adaptations, as has been detected in murine models^4, 10, 77^. Despite this tissue specificity, MΦs responded to *ex vivo* stimuli with similar transcriptional and functional profiles, concordant with responses in monocytes, though intestinal MΦs had a somewhat attenuated response in line with previous studies^51, 87^, possibly due to their maintenance in a site with a high burden of commensal microbes^87^. While MΦs are responsive to a wide variety of extrinsic signals^24, 25^, only some of which were evaluated here, these results suggest that tissue-specific and acute polarization programs are controlled by unique transcriptional regulators. Given the challenges with obtaining tissue resident MΦs from humans, we anticipate that identification of these stable signatures can also serve to aid recent efforts for *in vitro* establishment^26, 88, 89^ or maintenance^90^ of “tissue-adapted” MΦs.

As coordinators of innate immunity, dysregulation of MΦ function is central to many tissue-specific pathologies in humans, including inflammatory bowel disease, neuroinflammation, pulmonary fibrosis, and tumor growth^91–93^. The molecular and functional profiles revealed here in human monocytes and MΦs establish a homeostatic baseline from which we can better identify and understand tissue-specific dysregulations. While we focus here on *ex vivo* cytokine production among tissue-specified MΦs, our tissue signatures indicate that other functions—such as phagocytosis, efferocytosis, antigen presentation, and mobility—may mediate unique homeostatic and immune interactions for MΦs across the body. Our results can help define how tissue-adaptations affect diverse MΦ functions in different disease states and uncover molecular pathways with therapeutic potential.

## ACKNOWLEDGEMENTS

This work was supported by a Seed Networks for the Human Cell Atlas grant from the Chan Zuckerberg Initiative (CZF2019-002452) and NIH grants AI106697 awarded to P.A.Si. and D.L.F and AI057266 awarded to D.L.F. D.P.C. was supported by the Columbia University Graduate Training Program in Microbiology and Immunology (T32AI106711). P.A.Sz. was supported by a Canadian Institutes of Health Research (CIHR) Fellowship. Research reported here was performed in the Columbia Stem Cell Initiative Flow Cytometry Core, the Sulzberger Columbia Genome Center, and the Columbia Single Cell Analysis Core (supported by grant P30CA013696 and P30DK132710).

The content is solely the responsibility of the authors and does not necessarily represent the official views of the NIH. We wish to thank the donor families for their generosity and the exceptional efforts of the transplant coordinators and staff of LiveOnNY for making this study possible.

## AUTHOR CONTRIBUTIONS

D.P.C. designed and performed experiments, analyzed data, made figures, and wrote the manuscript. W.L.S. performed experiments, analyzed data, made figures, wrote the manuscript. D.C., S.B.W., and P.A.Sz. prepared samples for single cell sequencing. P.A.Si. and D.L.F. designed experiments, analyzed data, wrote, and edited the manuscript.

## DATA AVAILABILITY STATEMENT

CITE-seq data from the previously published human organ donor atlas is available at CellXGene: https://cellxgene.cziscience.com/collections/cc431242-35ea-41e1-a100-41e0dec2665b CITE-seq data generated in this study are available in NCBI GEO with the accession code GSE296669.

## SUPPLEMENTARY FIGURES

**Supplementary Figure 1:**
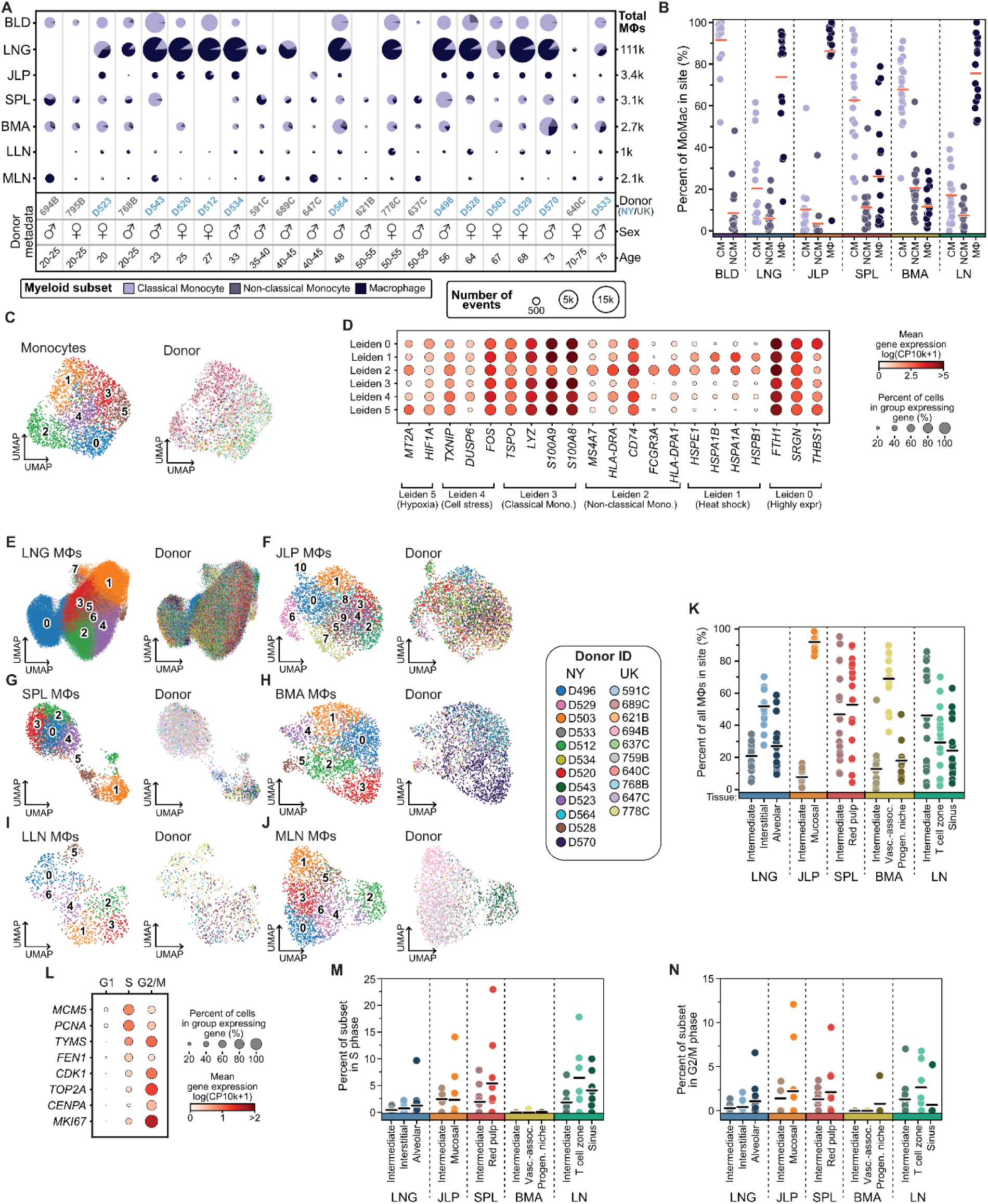
Clustering analysis reveals macrophage, but not monocyte, heterogeneity within each tissue site. **(A)** Dot plot of monocytes and macrophages (MΦs) extracted from an organ donor atlas of tissue immune cells. For each sample, pie charts depict cell type composition and dot size corresponds to number of events. **(B)** Strip plots quantifying the frequency of classical monocytes (CM), non-classical monocytes (NCM), and MΦs for each donor across each tissue site. Means are indicated by orange bars. Samples with fewer than 25 total cells were excluded. **(C)** Uniform manifold approximation & projection (UMAP) plots computed on a multimodal embedding of monocytes colored by Leiden cluster (left) or donor-of-origin (right). **(D)** Dot plot of selected markers associated with specific monocyte clusters. Dots are colored by mean gene expression (GEX) and size reflects the percent of cells expressing a gene. **(E-J)** UMAPs as in (C) but for MΦs isolated from lung (LNG), jejunum lamina propria (JLP), spleen (SPL), bone marrow (BMA), lung-associated lymph nodes (LLN), or mesenteric lymph nodes (MLN). **(K)** Strip plot as in (B) quantifying frequency of MΦ subsets as a proportion of total MΦs in each tissue site for each donor. **(L)** Dot plot as in (D) displaying expression of selected cell-cycle markers on MΦs isolated across tissues. MΦs are classified as either in G1, S, or G2/M phase of the cell cycle (see Methods). **(M-N)** Strip plots as in (B) quantifying the frequency of MΦs caught in cell cycle as a proportion of the total MΦ subset for each donor.

**Supplementary Figure 2:**
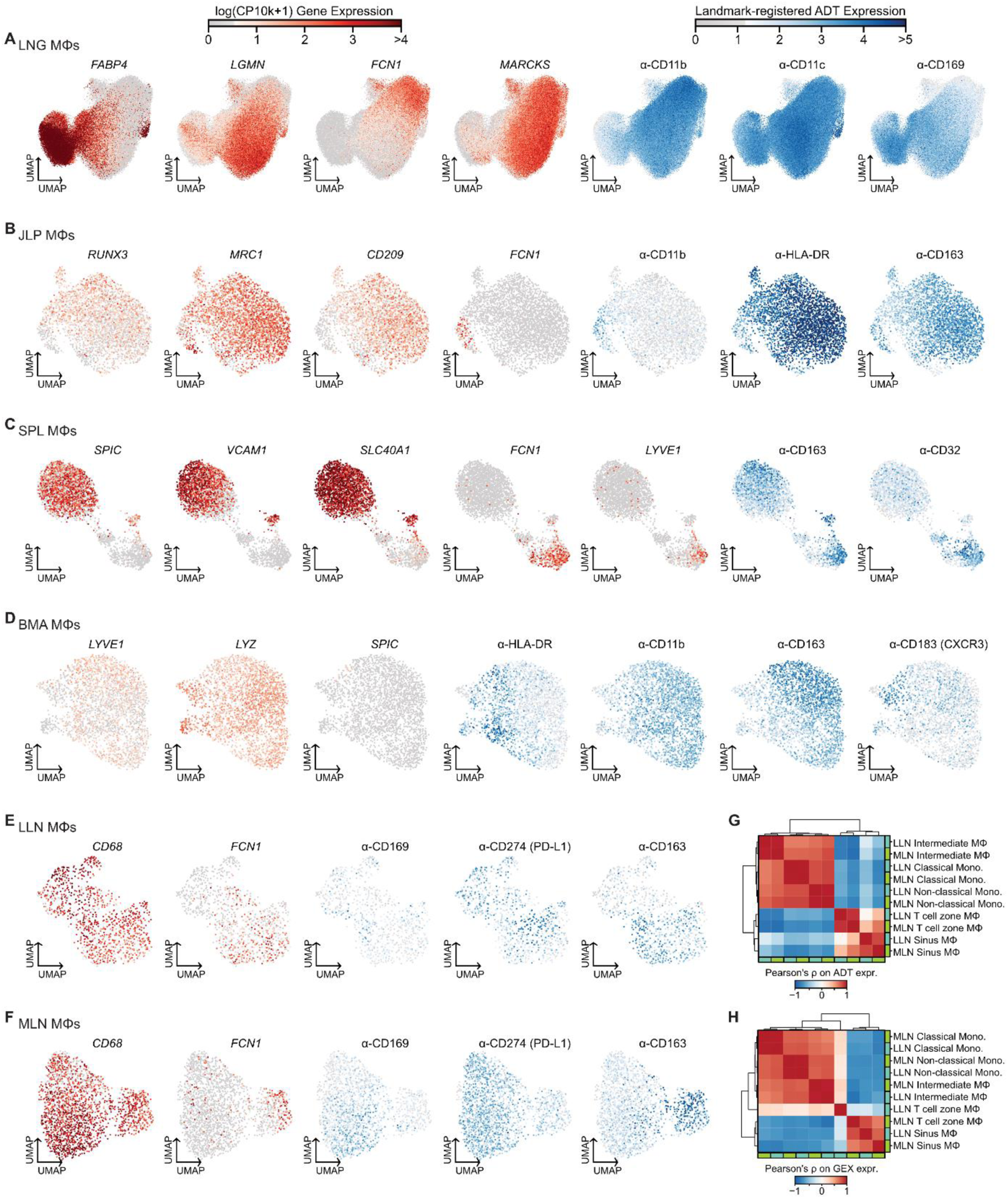
Gene and protein marker expression support macrophage subset delineation. **(A-F)** Uniform manifold approximation & projection (UMAP) plots computed on a multimodal embedding for macrophages (MΦs) isolated from lung (LNG; A), jejunum lamina propria (JLP; B), spleen (SPL; C), bone marrow (BMA; D), lung-associated lymph nodes (LLN; E), or mesenteric lymph nodes (MLN; F). UMAPs are colored by expression of selected markers, either gene expression (GEX; left, in reds), or landmark-registered antibody-derived tag (ADT) expression (right, in blues) **(G-H)** Clustered correlation matrices showing similarity between MΦ and monocyte subsets identified across the two lymph node sites. Pearson correlations were computed on a donor-integrated principal component analysis (PCA) embedding on normalized ADT (G) or gene (H) expression.

**Supplementary Figure 3:**
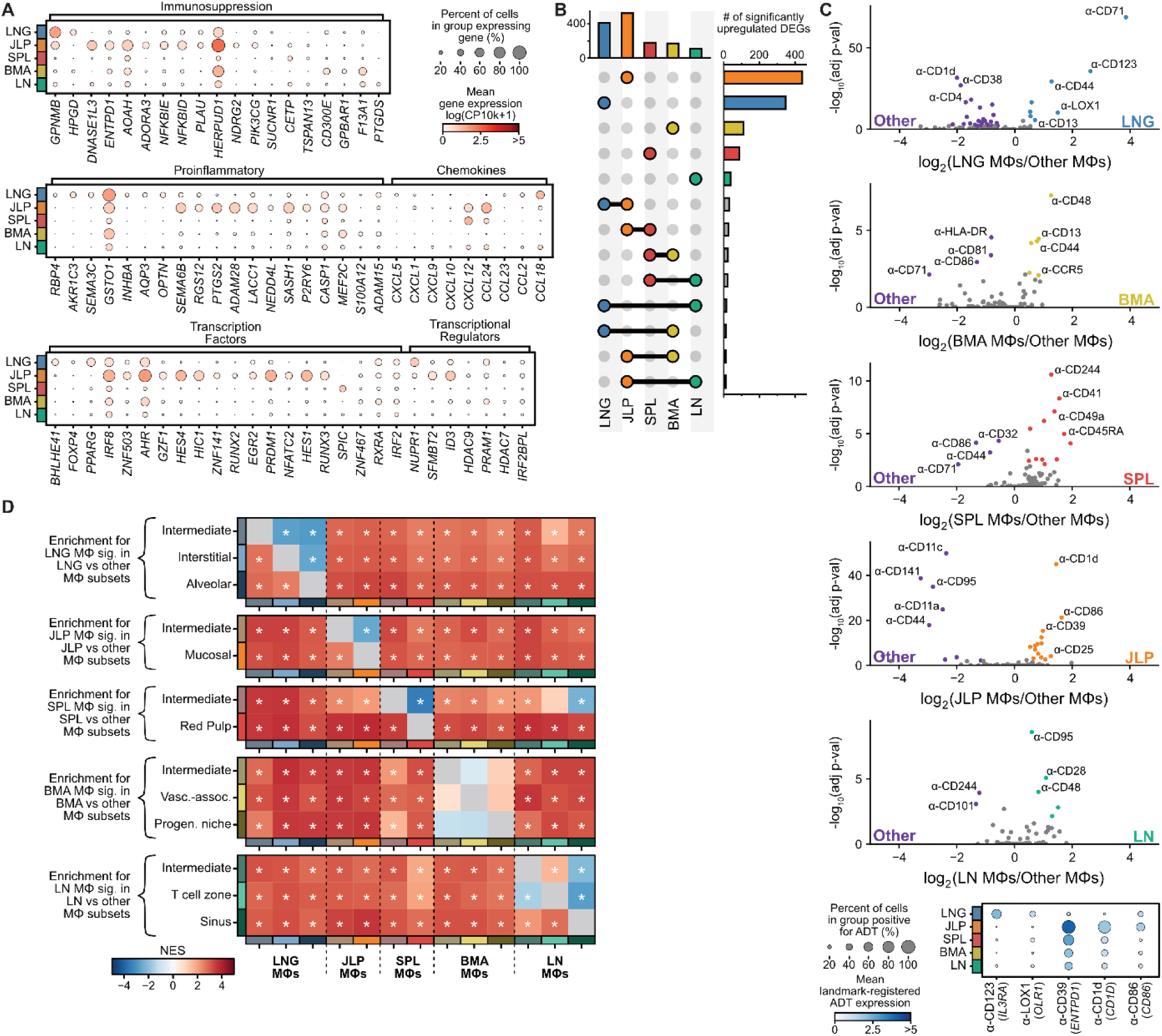
Human macrophages are tissue-adapted, with specific transcriptomes and surface proteomes. **(A)** Dot plots of selected differentially expressed genes (DEGs). Dots are colored by mean gene expression (GEX) and size reflects the percentage of cells expressing a gene. **(B)** UpSet plot quantifying overlap of significantly upregulated genes within each tissue-site. **(C)** Volcano plots (above) and dot plot (below) highlighting significantly differentially expressed antibody-derived tags (ADTs) by percent-positivity (adjusted p-value < 0.05 and |log_2_(fold-change)| > 0.5). Significant markers are highlighted in purple (downregulated) or a tissue-specific color (upregulated). Dot plots are colored by the mean landmark-registered ADT expression and size reflects the percent of MΦs with above-background expression for a surface marker **(D)** Heatmap displaying normalized enrichment score (NES) for each tissue signature when subsets within that tissue (rows) are compared to each other subset (columns). Asterisks denote significance (FDR adjusted p-value < 0.05) as assessed by pre-ranked gene set enrichment analysis (GSEA).

**Supplementary Figure 4:**
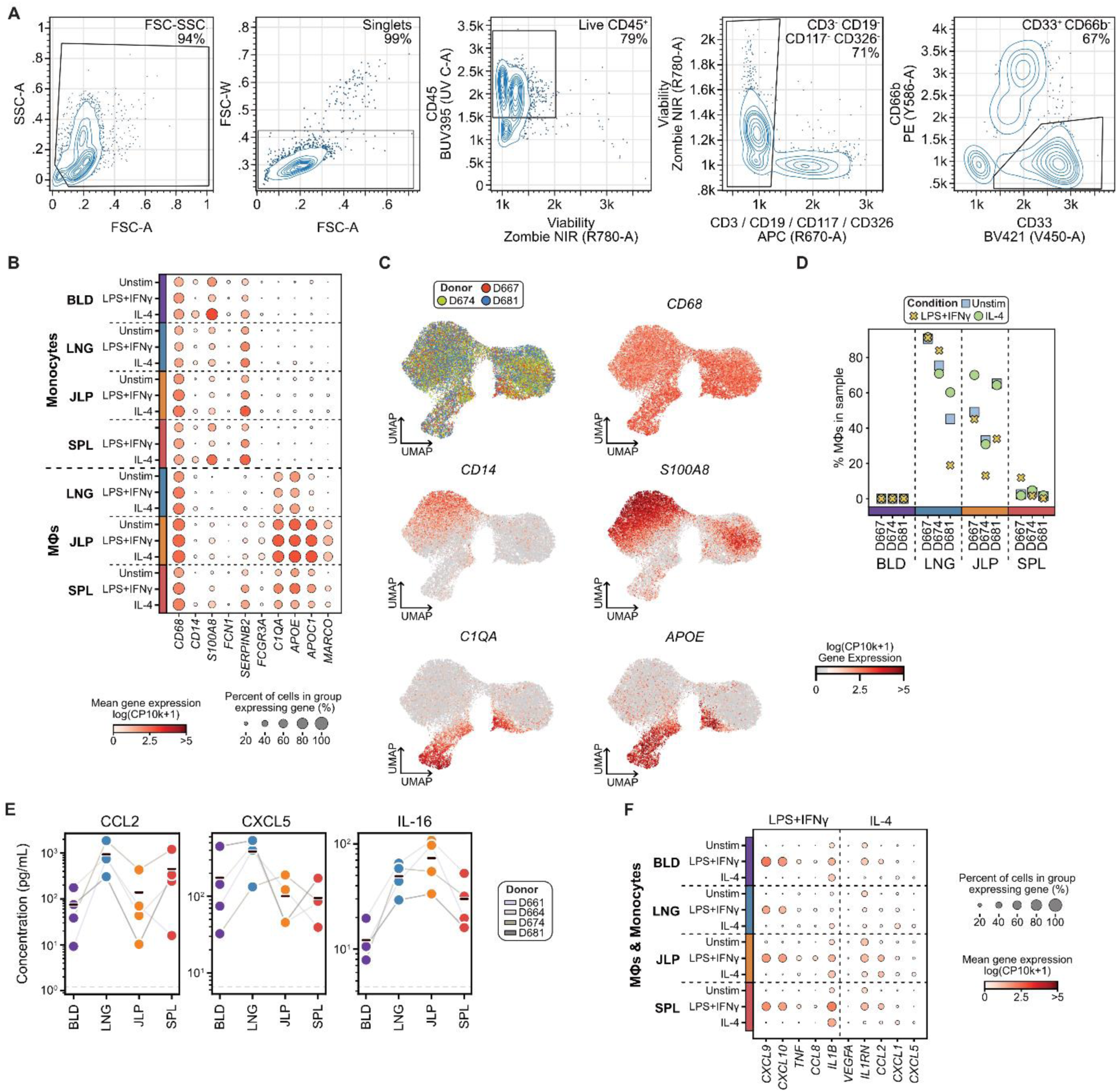
Sort, annotation, and differential expression of *ex vivo* stimulated monocytes and macrophages. **(A)** Example gating strategy for fluorescence-activated cell sorting (FACS) of macrophages (MΦs) and monocytes in a blood sample. **(B)** Dot plots of selected monocyte or MΦ cell type markers after annotation by unsupervised clustering. Dots are colored by mean gene expression (GEX) and size reflects the percentage of cells expressing a gene. **(C)** Uniform manifold approximation & projection (UMAP) plots colored by donor or GEX of selected markers **(D)** Strip plot displaying the frequency of MΦs recovered in each donor and tissue profiled by CITE-seq. Color/shape of points delineates treatment condition to display consistency of MΦ frequency within each sample. **(E)** Strip plots quantifying cytokine concentration in supernatants of unstimulated samples for analytes that align to tissue-specific gene expression results identified in Fig. 2. Means are indicated by black bars. Measurements from the same donor are connected by light gray lines. A dashed gray line displays the assay’s lower limit of detection. **(F)** Dot plots as in (B) displaying gene expression of selected cytokine markers.

